# Metagenomic sequencing of dung beetle intestinal contents directly detects and identifies mammalian fauna

**DOI:** 10.1101/074849

**Authors:** Conrad P.D.T. Gillett, Andrew J. Johnson, Iain Barr, Jiri Hulcr

**Affiliations:** School of Forest Resources and Conservation, University of Florida, 136 Newins-Ziegler Hall, Gainesville, Florida, 32611-0410, United States; School of Biological Sciences, University of East Anglia, Norwich, NR4 7TJ, United Kingdom

**Keywords:** Biodiversity measurement, eDNA, Environmental DNA, diet analysis, Illumina, mitochondrial, scarab beetles, Scarabaeidae, Scarabaeinae, Swaziland.

## Abstract

1. Cost, time, and expertise constraints limit traditional observation-based comprehensive biodiversity assessment. Therefore, surrogate focal taxa representative of wider biodiversity are commonly used as an imperfect ‘proxy’. Contemporary biodiversity assessments are also increasingly benefiting from the combination of high-throughput sequencing and metagenomic methodologies that enable identification of environmental DNA samples. However, there is a need for empirical studies combining the use of surrogate taxa with metagenomic approaches, that promise rapid and efficient biodiversity assessment.
2. We here tested for the first time the possibility of using the intestinal contents of wild-collected dung beetles (Scarabaeidae) as a source of mammalian DNA, in a metagenomics proof-of-concept approach to directly detect and identify mammals from an area of savanna-scrub in southern Africa. Dung beetles have been purveyed as an indirect proxy measure of mammalian diversity, owing to their dependence upon vertebrate dung as a food source, and the ease with which they can be comprehensively sampled using simple and repeatable trapping protocols, achievable much faster than vertebrate surveys.
3. Following shotgun sequencing of gut content DNA extractions from ten dung beetle species, we used *in silico* filters to identify mammals by searching the resulting reads against known mammalian mitochondrial DNA from online sequence repositories, matching 546 paired reads to known mitogenomes held in GenBank, and 634 reads to known mammal barcode sequences held in BOLD. Identified mammalian sequences were consistent with wild and domesticated ungulates known from the sampling site, and included blue wildebeest, plains zebra, and domestic cattle and goat. Four dung beetle samples yielded sufficient sequence data to successfully assemble the near-complete mitogenome of blue wildebeest at up to 21 X mean coverage, despite low initial DNA concentrations, unambiguously corroborating identification.
4. It is conceptually and practically possible to rapidly and economically apply metagenomic techniques in dung beetle gut sequencing to detect the presence of mammals upon whose dung the beetles have fed. Since the approach can be readily scaled up, it may prove to be of practical use as a complement to traditional biodiversity assessment methods, and should be tested in usefulness for detecting rare, endangered or cryptic mammal species.

## Introduction

Comprehensive and efficient biodiversity assessment is an important element of the decision making process leading to scientifically well-informed conservation policies (Reid et al. 1993; Humphries et al. 1995; Royal Society 2003). Amongst frequently assessed taxa, mammals are a charismatic group of important conservation value and concern, that provide key ecological services, and which are globally distributed (Schipper et al. 2008). However, they can be difficult to observe, and even more difficult to survey and monitor effectively because of their often reclusive habits, low abundances, small populations, and restricted distributions (Krebs 2006). Consequently, a great variety of field methods are used to monitor mammal biodiversity; most relying upon direct field observations of the animals (sightings, camera-trapping, sound recordings etc.) and their traces (droppings, footprints, burrows etc.) (Barnett 1995; Barnett & Dutton 1995; Krebs 2006). Such traditional biodiversity surveys are logistically complicated, dependent upon availability of taxonomic expertise, and require lengthy timescales to be comprehensive (Sutherland 2006). These limitations can pose challenges in detecting rare, endangered, secretive, nocturnal, or otherwise challenging taxa. It is therefore necessary to explore alternatives to traditional approaches, especially those making use of new technologies which could provide data of improved quality at lower effort and cost.

Contemporary metagenomics methodologies that take advantage of high throughput DNA sequencing are transforming the way biodiversity is measured (Bohmann et al. 2014; Corlett 2016). It is now possible to quantify animal diversity using DNA from a range of terrestrial, aquatic, and even airborne environmental samples (Creer et al. 2016). Moreover, this can be achieved with little or no prior taxonomic expertise, highlighting that metagenomic approaches can increasingly be purveyed as a potentially useful and cost-effective tool to overcome the “taxonomic impediment” (Yang et al. 2014). The contents of invertebrate intestines is becoming a widely-used source of environmental DNA (eDNA) in biodiversity sampling (Calvignac-Spencer et al. 2013a). That fragments of vertebrate DNA can be successfully extracted, amplified, sequenced, and identified from invertebrate digestive tracts has already been demonstrated in several studies, including those using blood-feeding leeches (Schnell et al. 2012) and mosquitos (Towzen, Brower & Judd 2008; Mehus & Vaughan 2013), and carrion-feeding flies (Calvignac-Spencer et al. 2013b) as the source material.

Dung beetles (Coleoptera: Scarabaeinae) are a diverse group of insects (c. 6000 described species) occurring in a great variety of ecozones and on all continents except Antarctica, attaining their greatest diversity in the Afrotropical region (Cambefort 1991). They have hitherto been routinely used as an *indirect* surrogate for mammalian diversity (Gardner et al. 2008), primarily owing to their reliance upon vertebrate dung and carrion for nutrition, resulting in a correlation in their diversities. Dung beetles can also be efficiently and economically sampled (Halffter & Favila 1993; Spector 2006; Gardner et al. 2008; Larsen 2011) using both active (baited) and passive (not reliant upon a bait lure) trapping protocols. Because dung beetles contain both generalist and specific coprophagous and necrophagous species (Halffter & Mattthews 1966; Hanski & Cambefort 1991; Larsen, Lopera & Forsyth 2006), they are potentially a source of a great diversity of vertebrate DNA, including that of cryptic smaller species not easily surveyed through direct visual observation or physical trapping. Yet, they have not hitherto been used as a potential source of vertebrate DNA to *directly* sample vertebrate diversity, despite their gut contents likely containing persisting fragmented DNA originating from shed gut epithelial and blood cells from the vertebrate source of the dung, voided in the dung, and consumed by the beetles.

Previous studies using invertebrates as a sampling tool for vertebrate DNA have primarily relied upon PCR-amplification and Sanger sequencing of a few mitochondrial genes, followed by subsequent taxonomic assignment using BLAST searches (Altschul et al. 1990) against known vertebrate sequences (Calvignac-Spencer et al. 2013a; Drummond et al. 2015). The present study aimed to use modern high-throughput metagenomics methodology to test the feasibility of *directly* identifying vertebrate mitochondrial DNA (mtDNA) *in silico* from dung beetle intestinal content DNA extractions as a proof-of-concept, and to discuss the potential applications of this technique in biodiversity assessment. In particular, we wanted to circumvent bias-prone and time- and resource-intensive PCR amplification steps, through direct ‘shotgun’ sequencing of the beetle gut extractions. mtDNA was selected as the focal target molecule, both because of its widespread use in species-level identification of animal taxa (and the corresponding availability of identified reference sequences on publicly accessible repositories), and because of the increased chance of sequencing success as it is present in multiple copies in animal cells. In line with these goals, the objectives were to answer the following questions: A) How much vertebrate mtDNA can be detected from the dung beetle gut contents? B) Can this mtDNA be used for identification of mammal taxa? C) Does the quality and quantity of resulting reads allow for assembly of longer mitogenomic contigs? D) If any vertebrate taxa are identified, are they consistent with the known fauna of the sampled area?

## Materials and methods

### Specimen sampling

Dung beetles were sampled on the 25^th^ and 26^th^ of March 2016 at the Mbuluzi Game Reserve in the Lubombo Region of eastern Swaziland (approx. 26.1°S; 32.0°E; 200 masl), an area predominantly covered in savanna-forest and scrub vegetation. This managed reserve is home to a wide diversity of mammals (Appendix S1) and other vertebrates, including several species of ungulates, together with a correspondingly rich fauna of scarabaeine dung beetles. Immediately prior to, and during the course of sampling, the area experienced considerable rainfalls following a prolonged period of drought, leading to conspicuous dung beetle activity. The beetles were collected passively (i.e. not attracted to a dung bait) using two flight interception traps (FIT) set in typical savanna-forest. Each FIT consisted of a 1.5 m X 1.0 m fine nylon mesh sheet, held taut between the trunks of two shrubs, with the lower edge suspended approximately 15 cm above ground level. Several plastic trays, of approximately 10 cm depth, and half-filled with water, were placed on the ground immediately below and in line with the mesh, to collect beetles intercepted in flight. The traps were inspected twice daily over the two days, and collected specimens were preserved individually in plastic tubes containing 96% ethanol.

The collected specimens were identified morphologically using a variety of pertinent taxonomic literature (Ferreira 1972; Palestrini 1992; Davis, Frolov & Scholtz 2008; Deschodt, Davis & Scholtz 2015; Pokorny & Zídek 2015) and through comparison to specimens in the first author’s entomological reference collection. From among them, ten species representing a wide phylogenetic, ecological, and size diversity, were selected for intestinal content DNA extraction.

The selected specimens were dissected individually under a fume hood, using sharp scalpels and fine forceps, adhering to standard aseptic techniques to minimise environmental contamination. The elytra were either raised or removed to expose the dorsal abdominal tergites. An incision was made along the longitudinal axis of the abdomen and as much of the intestine and its contents was removed as possible. In most cases it was obvious that the gut contained faecal matter, but in cases where it appeared empty, the specimen was rejected, and another of the same species was selected for dissection. In addition to the individual gut samples, one pooled sample of the dung adhering to the beetle specimens was prepared from the suspension in the preserving ethanol, to investigate whether this residual dung source contained detectable mammal mtDNA. The ethanol-dung suspension from each preserved beetle was briefly centrifuged at 10,000 RPM, to separate out the bulk of the dung in the bottom of the centrifuge tube. Approximately 100 μl of this dung-ethanol mixture from each specimen was pooled together in a separate tube and mixed on a vortex mixer. 100 μl of this resulting pooled suspension was used in the subsequent DNA extraction.

### DNA extraction, library preparation and sequencing

DNA was extracted from each intestinal dissection sample using Qiagen DNeasy blood & tissue spin column kits (Qiagen). The resulting DNA concentrations and purities were quantified independently on both a Nanodrop spectrophotometer and a Qubit fluorometer (Thermo-Fisher Scientific), using a dsDNA high-sensitivity assay kit (Invitrogen). An individual and unique sequencing library preparation was constructed for each DNA sample, using a NEBNext Ultra DNA Library Prep Kit (New England BioLabs) for Illumina, with a targeted mean insert size of 500 bp, and entirely unique dual indexes. Libraries were further size selected using the SageELF electrophoresis system (Sage science), and subsequently pooled at equimolar concentrations and sequenced on a single run of a NextSeq sequencer (Illumina) with high output 150 bp paired-end reads. The sequencing run was shared with other insect RNAseq and DNA libraries (none associated with mammals), such that each of the eleven dung beetle libraries was allocated 1/25 of the sequencing run. Sequencing was undertaken at the University of Florida Interdisciplinary Center for Biotechnology.

## Sequence analysis

### Quality control

Prior to mammalian sequence identification, all the raw reads were filtered to remove low quality reads or remaining adapter sequences using Trimmomatic (Bolger et al. 2014). Both the trimmed forward (R1) and reverse (R2) orientation reads were used in separate *in silico* search strategies against two sequence repository databases, in order to identify mammalian mtDNA matching to the reads. The first incorporated searches against mammalian mitochondrial genomes (mitogenomes) retrieved from GenBank (Benson et al. 2013), and the second incorporated searches against all mammalian sequences held in the Barcode of Life Data System (BOLD) database (Ratnasingham et al. 2007). Additionaly, *de novo* assembly of reads matching to the mammalian mitogenomes was undertaken, followed by taxonomic assignment of any resulting contigs using BLAST searches against GenBank, as detailed below. Bioinformatics analyses were undertaken using the University of Florida HiPerGator 2.0 supercomputer. Figure 1 summarises the main workflow steps undertaken for these analyses.

**Figure 1:**
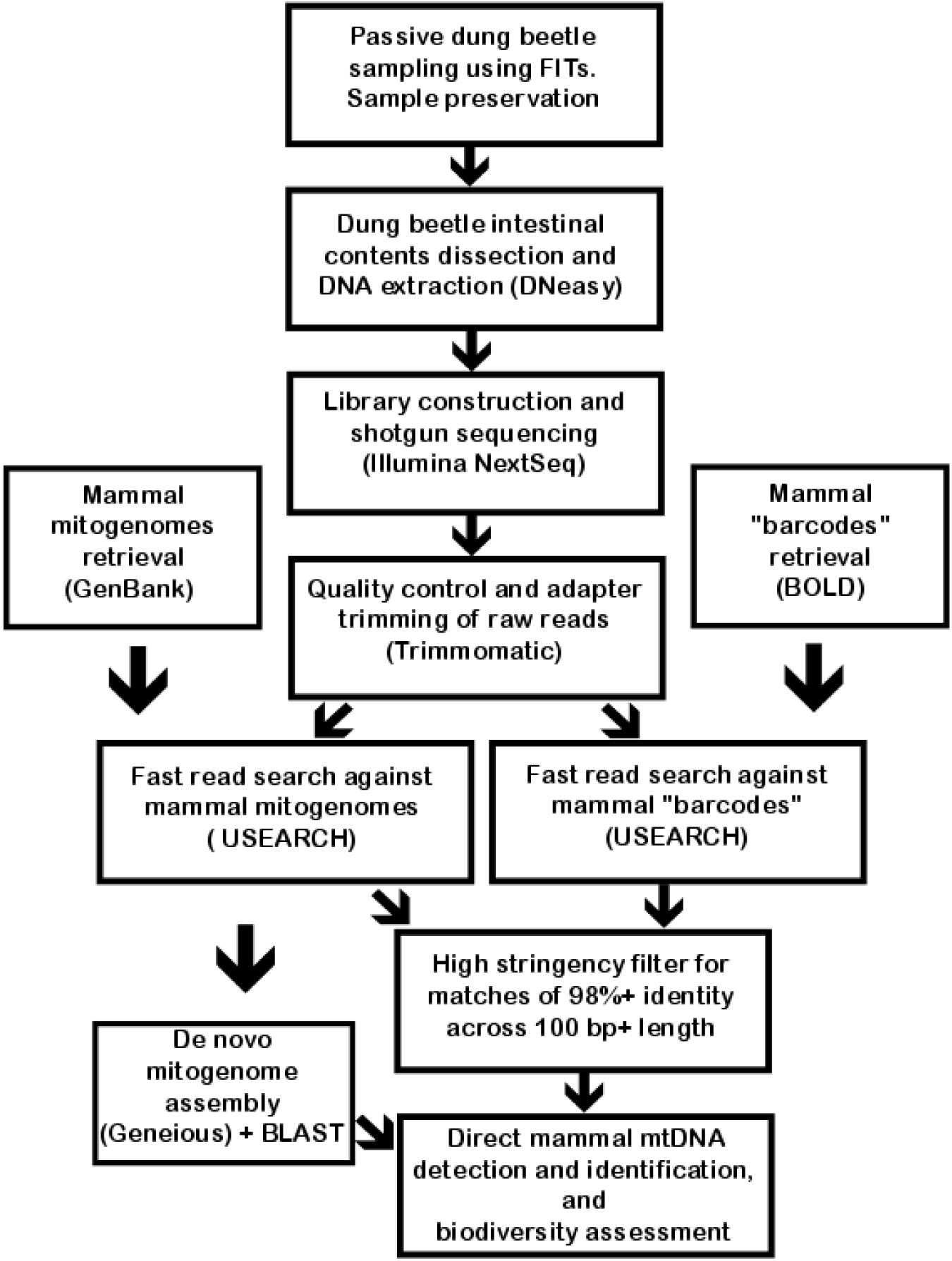
Schematic workflow of steps undertaken in the dung beetle gut DNA extraction metagenomics analysis.

### Mammal mitogenome search

To retrieve mammal mtDNA from the sequence pool, we matched the sequence output against a custom FASTA format database of complete and near-complete mammal mitogenomes retrieved from GenBank, representing 25 diverse species (Appendices S2 and S3). This included eight wild ungulate species, and two primate species known to occur at the sampling site, in addition to five domestic animal species, nine other (mostly African) mammals, and a human mitogenome. Initial low-stringency searches were undertaken using all the reads from each sample separately against the mitogenome database, to filter for mammalian-like sequences. Searches were undertaken using the USEARCH global alignment search algorithm (Edgar 2010), retaining only each read’s closest match (i.e. the top ‘hit’) to a mammal mitogenomic sequence, with a minimum of 90% sequence identity. USEARCH was used because it has been empirically shown to offer orders of magnitude faster searching than BLAST in practical applications (Edgar 2010). The matched reads were thereafter filtered, recording only those where both corresponding R1 and R2 paired-reads matched the same reference mammal mitogenome, each with a stringency of 98% or higher sequence identity and a minimum of 100 bp coverage, ensuring that a highly conservative level of sequence identification was employed.

### Mammal mitogenome assembly

Mitogenome-based identification of mammal species was achieved for each sample separately, through the *de novo* assembly of all the reads (in both R1 and R2 orientations) matching to a mammal mitogenome reference with 90% or greater identity, i.e. using the retained reads following the initial search using USEARCH, as detailed above. Assembly was undertaken with Geneious (Kearse et al. 2012) using the native high sensitivity settings (15% maximum gaps per read, maximum gap size 2bp, minimum overlap 25bp, minimum overlap identity 80%, maximum mismatches per read 30%, maximum ambiguity 4). The resulting consensus sequence of each of the longest assembled mitogenomic contigs per sample were used in a BLAST search against GenBank to determine sequence identity.

### Mammal DNA “barcode” search

All mammalian barcode sequences held on the BOLD database were retrieved using the search term “Mammalia” on the Public Data Portal retrieval interface, and downloaded as a reference database in a single FASTA file on the 28^th^ July 2016 (Appendix S4). This database contained not only *cox1* sequences (often considered the standard “barcode” for animals) but also sequences from several other mitochondrial genes. Searches were undertaken using all the reads from each sample separately against the barcode database using USEARCH, and retaining only each read’s closest match to a mammal sequence, with a minimum of 98% identity. The matched reads were thereafter filtered, recording only those where either R1 or R2 reads matched a mammal sequence with a stringency of 98% or higher sequence identity across a minimum sequence length of 100 bp., ensuring a highly conservative level of sequence identification was employed.

## Results

### Beetle identification, DNA extraction and gut-content sequencing

The ten species and samples of dung beetle selected for dissection and gut-content sequencing are listed in Appendix S5. They belong to ten different genera and to eight tribes of Scarabaeinae, including representatives of the two major ecological groups, the “tunnelers” and the “rollers”, as defined by Cambefort & Hanski (1991). The DNA extraction concentrations varied greatly, between 1.44 – 456.85 ng/ μl as measured by Nanodrop, and between < 0.1 – 74.0 ng/μl as measured by Qubit (Appendix S5). DNA concentration was broadly correlated to beetle size, with extractions from smaller beetles generally resulting in lower concentrations (data not shown). Although two samples (DB007 and DB009) yielded very low DNA concentrations, almost undetectable on the Qubit, their library preparations and sequencing was nevertheless undertaken successfully.

A total of 191,888,578 paired sequence reads were obtained for all 11 sample libraries combined, with individual sample libraries producing between 8,450,155 (DB007) and 25,459,419 (DB001) paired reads. Following adapter trimming for each sample, the percentage of reads surviving quality control in both directions varied between 79.1% (DB010) and 99.88% (DB006). The percentage of dropped reads for each sample varied between 0.01% (DB006) and 0.82% (DB002) (Appendix S6).

## Sequence identification

### Mammal mitogenome search

546 post-quality-controlled paired sequence reads obtained from nine of the ten dung beetle gut extractions (all except DB002), as well as the pooled dung sample, successfully matched mammalian mitogenomic sequences. Seven species of mammals accounted for all the matches, with the majority of matches (481 or 88%) to Blue wildebeest (*Connochaetes taurinus*), which was detected in eight of the ten dung beetle gut extractions, in addition to the pooled dung sample. Other mammal species identified, and known to occur within or in the environs of Mbuluzi, included Plains Zebra (*Equus quagga*, 21 matches), Domestic cattle (*Bos taurus*, nine matches) and Domestic goat (*Capra hircus*, eight matches). A single pair of reads matched to Blesbok (*Damaliscus pygargus*), a species not known to be present at Mbuluzi. Because this was unexpected, the two corresponding reads matching to blesbok were used in a BLAST search against all sequences on GenBank, which revealed a top match of 99% identity (150 of 151 bp matching) to blue wildebeest (GenBank accession JN632627) for the R1 orientation, and a match of 100% identity (across all 151 bp) to domestic goat (KR349363) for the R2 orientation. We therefore reject the identification of blesbok, and cannot distinguish between the two other species based on available information. A total of ten paired reads matched to domestic mouse (*Mus musculus*) sequences, and 23 paired reads matched human mitogenomic sequences. Sample DB004 (*Onitis aeruginosus*) yielded the highest number of mammalian matches (145), and only the extraction from sample DB002 (*Anachalcos convexus*) did not yield any matches. The pooled dung sample resulted in 16 mammalian matches to three species: blue wildebeest, plains zebra, and human. Table 1 summarises the number of matches to mammal mitogenome sequences from each of the ten dung beetle gut extractions, and the pooled sample.

**Table 1:** Paired sequence read matches to 25 mammal mitogenomes retrieved from GenBank, with 98% minimum identity over a minimum of 100 bp.

| Common name | Scientific name | GenBank accession code | DB001 | DB002 | DB003 | DB004 | DB005 | DB006 | DB007 | DB008 | DB009 | DB010 | DB011 | Total |
| --- | --- | --- | --- | --- | --- | --- | --- | --- | --- | --- | --- | --- | --- | --- |
| Domestic cattle | *Bos taurus* | EU177832 | 1 | 0 | 0 | 2 | 2 | 0 | 2 | 1 | 1 | 0 | 0 | 9 |
| Blesbok | *Damaliscus pygargus* | FJ207530 | 0 | 0 | 0 | 0 | 1 | 0 | 0 | 0 | 0 | 0 | 0 | 1 |
| Domestic goat | *Capra hircus* | GU295658 | 0 | 0 | 0 | 1 | 2 | 0 | 1 | 2 | 2 | 0 | 0 | 8 |
| Blue wildebeest | *Connochaetes taurinus* | JN632627 | 1 | 0 | 43 | 142 | 65 | 36 | 40 | 40 | 105 | 0 | 9 | 481 |
| Plains Zebra | *Equus quagga* | JX312733 | 0 | 0 | 0 | 0 | 0 | 0 | 0 | 0 | 1 | 19 | 1 | 21 |
| House mouse | *Mus musculus* | NC_005089 | 2 | 0 | 0 | 0 | 0 | 0 | 0 | 1 | 0 | 0 | 0 | 3 |
| Human | *Homo sapiens* | NC_012920 | 1 | 0 | 0 | 0 | 0 | 0 | 9 | 5 | 2 | 0 | 6 | 23 |
| Total |  |  | 5 | 0 | 43 | 145 | 70 | 36 | 52 | 49 | 111 | 19 | 16 | 546 |

### Mammal mitogenome assembly

The near complete mitochondrial genome (> 16,400 bp) of blue wildebeest was assembled from the reads in four of the dung beetle gut extractions (DB003, DB004, DB005, and DB009). These assemblies matched the same blue wildebeest mitogenome sequence on GenBank (JN632627) with an E-value of 0, and with > 99% identity across their full assembly lengths. Mean coverage depths for these complete four assemblies varied between 7.3 X (standard deviation = 3.0) in sample DB003, and 21.8 X (standard deviation = 5.6) in sample DB004. The longest mitogenomic assemblies from three additional samples (DB006-008), varying in length between 2668-5797 bp, also matched the same blue wildebeest sequence on GenBank, each with an E-value of 0 (coverage varying between 4.7-7.4 X). Sample DB010 resulted in a short 350 bp assembly (of three read pairs) which matched a plains zebra mitogenome sequence on GenBank (JX312729) with an E-value of 6×10^−176^ and 99% identity across the entire assembly length. Other short assemblies included a 313 bp assembly matching a house mouse mitogenome (NC_005089) with an E-value of 3×10^−154^and 98% identity across the entire assembly (sample DB001), and an 837 bp assembly matching a human mitogenome (KX495641) with an E-value of 0 and 91% identity across the entire assembly (sample DB011). Sample DB002 resulted in a single very short assembly of 88 bp, which could not be significantly matched to any sequence on GenBank, and is not considered further in this study. Table 2 summarises information on the longest mitogenome assemblies obtained for each of the dung beetle gut extractions and their taxonomic assignments using BLAST.

**Table 2:**
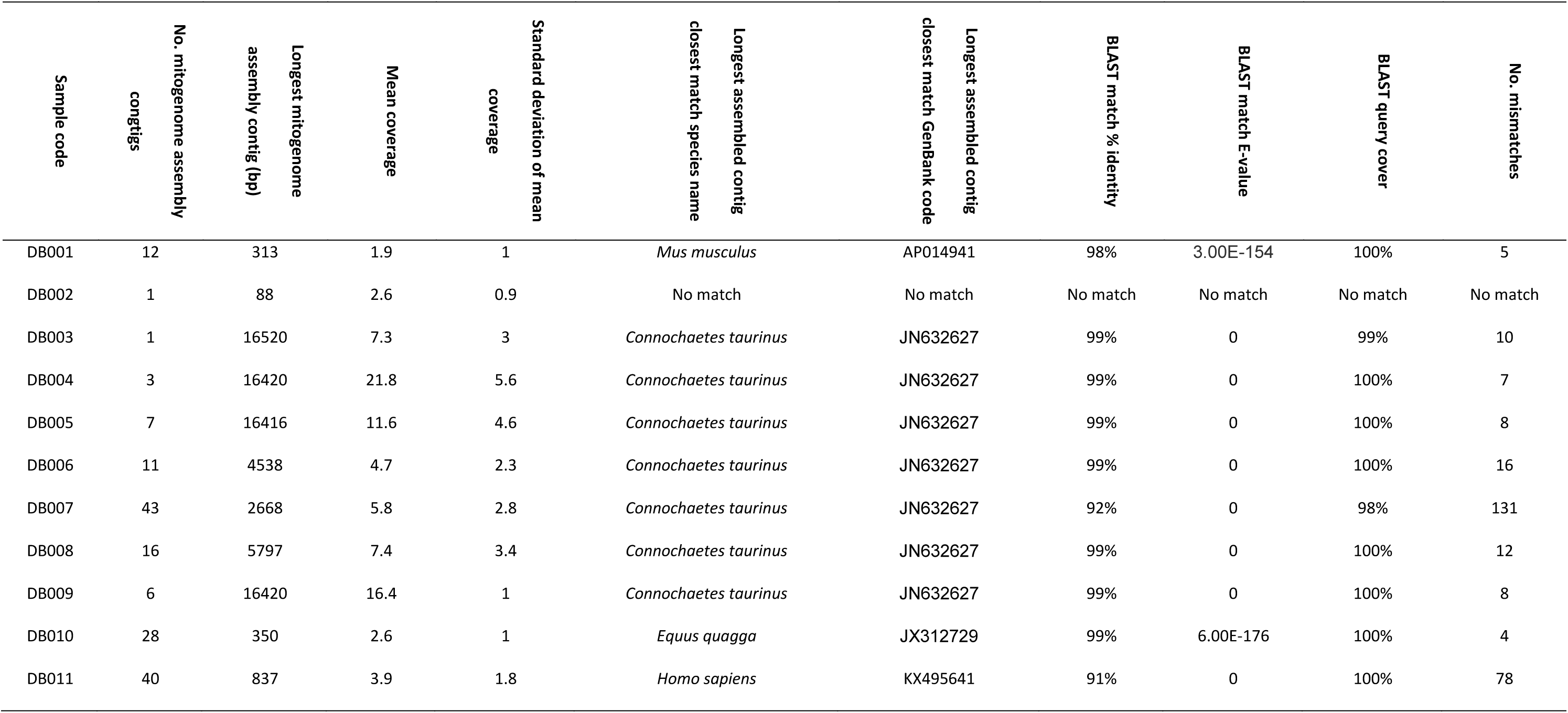
Summary of the longest *de novo* mitogenome assembly contigs obtained for each of the dung beetle gut extractions and their taxonomic assignments using BLAST.

### Mammal “barcode” search

The downloaded BOLD reference database comprised of 67,779 sequences, representing 1931 named species of mammals. Following the search of this database against the reads from each sample, a total of 634 reads matched mammalian sequences belonging to seven species. All samples except DB002 resulted in matches to mammal sequences in BOLD. The barcode analysis corroborated the mitogenomic analysis: the majority of matches (492 or 77%) were to four blue wildebeest sequences (488 matches) and to one sequence identified only to the genus *Connochaetes* (four matches). Blue wildebeest matches were found in all samples except DB001, DB0002, and DB010. Two reads from sample DB010 matched a sequence from plains zebra. A total of 22 matches to eight different sequences belonging to domestic cattle were recorded, including one sequence most closely matching to the zebu subspecies *B. taurus indicus*. Two reads from sample DB001 matched to a house mouse sequence, and 56 reads from seven samples (all except DB001, DB002, DB006, and DB010) matched to three sequences identified only as ‘Mammalia’. A total of 46 reads from seven of the samples (all except DB001, DB002, DB004, and DB006) matched to twelve different human sequences, with the majority (29 or 63%) matching to one sequence (BOLD accession CYTC1123-12). Additionally, five reads from three samples (DB004, DB005, and DB008) matched to two 16S rDNA sequences from water buffalo (*Bubalus bubalis*), and nine reads from four samples (DB004, DB005, DB007, and DB009) matched to one mtDNA control region D-loop sequence from white-tailed deer (*Odocoileus virginianus*). The latter two species of mammals are not known from Mbuluzi, therefore these matching reads were investigated further. Following BLAST searches of these reads against GenBank, it was found that the reads originally matching water buffalo sequences in BOLD, matched to blue wildebeest, waterbuck (*Kobus ellipsiprymnus*), Hunter’s antelope (*Beatragus hunteri*), black wildebeest (*C. gnou*), common duiker (*Silvicapra grimmia*), and southern reedbuck (*Redunca arundinum*) sequences in GenBank with 100% sequence identity and coverage. Similarly, the reads originally matching white-tailed deer in BOLD, matched to blue wildebeest and domestic goat sequences in GenBank with 100% sequence identity and coverage.

Table 3 indicates, for each of the samples, the number of reads matching to a mammalian sequence in the BOLD database with 98% or greater identity over at least 100 bp.

**Table 3:** The number of R1 or R2 reads, for each dung beetle gut extraction sample, matching to a mammalian sequence in the BOLD database with 98% or greater identity over at least 100 bp.

| Common<br>name | Scientific<br>name | BOLD<br>accession<br>code | DB001 | DB002 | DB003 | DB004 | DB005 | DB006 | DB007 | DB008 | DB009 | DB010 | DB011 | Total |
| --- | --- | --- | --- | --- | --- | --- | --- | --- | --- | --- | --- | --- | --- | --- |
| Domestic cattle | *Bos taurus* | CYTC5225-12 | 2 | 0 | 2 | 2 | 0 | 0 | 0 | 1 | 2 | 0 | 0 | 9 |
| Domestic cattle | *Bos taurus* | CYTC422-12 | 0 | 0 | 0 | 0 | 0 | 0 | 0 | 2 | 0 | 0 | 0 | 2 |
| Domestic cattle | *Bos taurus* | CYTC427-12 | 0 | 0 | 0 | 0 | 0 | 0 | 0 | 4 | 1 | 0 | 0 | 5 |
| Domestic cattle | *Bos taurus* | GBMA0356-06 | 0 | 0 | 0 | 0 | 1 | 0 | 0 | 0 | 0 | 0 | 0 | 1 |
| Domestic cattle | *Bos taurus* | GBMA0411-06 | 0 | 0 | 0 | 0 | 1 | 0 | 0 | 0 | 0 | 0 | 0 | 1 |
| Domestic cattle | *Bos taurus* | GBMA2505-09 | 0 | 0 | 0 | 0 | 1 | 0 | 0 | 0 | 0 | 0 | 0 | 1 |
| Domestic cattle | *Bos taurus* | GBMA9027-15 | 0 | 0 | 0 | 0 | 1 | 0 | 1 | 0 | 0 | 0 | 0 | 2 |
| Domestic cattle<br>(zebu) | *Bos taurus indicus* | CYTC5193-12 | 0 | 0 | 0 | 0 | 1 | 0 | 0 | 0 | 0 | 0 | 0 | 1 |
| House mouse | *Mus musculus* | CYTC662-12 | 2 | 0 | 0 | 0 | 0 | 0 | 0 | 0 | 0 | 0 | 0 | 2 |
| Blue<br>wildebeest | *Connochaetes taurinus* | CYTC1531-12 | 0 | 0 | 46 | 158 | 70 | 14 | 37 | 32 | 78 | 0 | 3 | 438 |
| Blue<br>wildebeest | *Connochaetes taurinus* | GBMA4117-12 | 0 | 0 | 4 | 12 | 8 | 3 | 6 | 6 | 8 | 0 | 1 | 48 |
| Blue<br>wildebeest | *Connochaetes taurinus* | GBMA5650-13 | 0 | 0 | 0 | 1 | 0 | 0 | 0 | 0 | 0 | 0 | 0 | 1 |
| Blue<br>wildebeest | *Connochaetes taurinus* | GBMA4150-12 | 0 | 0 | 0 | 1 | 0 | 0 | 0 | 0 | 0 | 0 | 0 | 1 |
| Wildebeest<br>(genus) | *Connochaetes* | GBMA5924-13 | 0 | 0 | 1 | 1 | 0 | 0 | 0 | 2 | 0 | 0 | 0 | 4 |
| Plains zebra | *Equus quagga* | GBMA10041-15 | 0 | 0 | 0 | 0 | 0 | 0 | 0 | 0 | 0 | 2 | 0 | 2 |
| Mammal | *Mammalia* | GBMA5729-13 | 0 | 0 | 0 | 1 | 0 | 0 | 0 | 0 | 0 | 0 | 0 | 1 |
| Mammal | *Mammalia* | GBMA5771-13 | 0 | 0 | 0 | 1 | 0 | 0 | 0 | 0 | 1 | 0 | 0 | 2 |
| Mammal | *Mammalia* | GBMA5776-13 | 0 | 0 | 11 | 14 | 6 | 0 | 2 | 6 | 13 | 0 | 1 | 53 |
| Human | *Homo sapiens* | CYTC1123-12 | 0 | 0 | 0 | 0 | 0 | 0 | 20 | 1 | 2 | 0 | 6 | 29 |
| Human | *Homo sapiens* | CYTC1939-12 | 0 | 0 | 0 | 0 | 0 | 0 | 0 | 0 | 0 | 1 | 0 | 1 |
| Human | *Homo sapiens* | CYTC173-12 | 0 | 0 | 0 | 0 | 0 | 0 | 1 | 0 | 0 | 0 | 0 | 1 |
| Human | *Homo sapiens* | CYTC3074-12 | 0 | 0 | 0 | 0 | 0 | 0 | 1 | 0 | 0 | 0 | 0 | 1 |
| Human | *Homo sapiens* | CYTC3232-12 | 0 | 0 | 0 | 0 | 0 | 0 | 2 | 0 | 0 | 0 | 0 | 2 |
| Human | *Homo sapiens* | CYTC153-12 | 0 | 0 | 1 | 0 | 0 | 0 | 0 | 0 | 0 | 0 | 0 | 1 |
| Human | *Homo sapiens* | CYTC2122-12 | 0 | 0 | 0 | 0 | 0 | 0 | 1 | 0 | 0 | 0 | 0 | 1 |
| Human | *Homo sapiens* | CYTC2553-12 | 0 | 0 | 0 | 0 | 0 | 0 | 1 | 0 | 0 | 0 | 0 | 1 |
| Human | *Homo sapiens* | CYTC2203-12 | 0 | 0 | 0 | 0 | 0 | 0 | 0 | 0 | 0 | 0 | 1 | 1 |
| Human | *Homo sapiens* | GBHS5339-09 | 0 | 0 | 0 | 0 | 2 | 0 | 2 | 0 | 2 | 0 | 0 | 6 |
| Human | *Homo sapiens* | GBMA12205-15 | 0 | 0 | 0 | 0 | 0 | 0 | 1 | 0 | 0 | 0 | 0 | 1 |
| Human | *Homo sapiens* | GBMA10457-15 | 0 | 0 | 0 | 0 | 0 | 0 | 1 | 0 | 0 | 0 | 0 | 1 |
| Water buffalo | *Bubalus bubalis* | YAKJC023-13 | 0 | 0 | 0 | 1 | 0 | 0 | 0 | 1 | 0 | 0 | 0 | 2 |
| Water buffalo | *Bubalus bubalis* | YAKJC010-13 | 0 | 0 | 0 | 1 | 1 | 0 | 0 | 0 | 1 | 0 | 0 | 3 |
| White-tailed<br>deer | *Odocoileus virginianus* | CYTC474-12 | 0 | 0 | 0 | 2 | 1 | 0 | 2 | 0 | 4 | 0 | 0 | 9 |
| Total |  |  | 4 | 0 | 65 | 195 | 93 | 17 | 78 | 55 | 112 | 3 | 12 | 634 |

## Discussion

Mammalian mtDNA can be successfully extracted, sequenced and identified from the intestinal contents of dung beetles without targeted PCR-amplification. This now establishes dung beetles as a useful source of mammalian DNA that can be directly identified, and not only a useful focal taxon for indirectly estimating wider biodiversity. Mammalian DNA sampling via dung beetles is highly scalable: whilst dung beetles are easily collected in their thousands, even a small number of beetle specimens (10 in this study) was sufficient to identify several of the common mammals present at the sampling site. The majority (90%) of our gut extraction samples resulted in sequences assignable to known mammalian mtDNA, whilst 60% contained DNA from more than one species of mammal, demonstrating the effectiveness of sampling. Results based both upon searches against mammal mitogenomes and barcodes were highly congruent, corroborated each other, demonstrated repeatability, and indicated that sufficient sequence data is generated without the need for locus-specific PCR amplification, with its associated bias (Beng et al. 2016).

Both mitogenome- and barcode-matching strategies identified nearly identical sets of mammals, displaying a similar distribution in their proportion of matching reads. Blue wildebeest, zebra, domestic cattle and goat, and humans were the source of most of the assignable sequences, with the majority of sequences (88% in the mitogenome search, and 77% in the BOLD search) matching to blue wildebeest, a common grazing ungulate at the sampling site.

Three species of mammal not known to occur at Mbuluzi were also identified via a small number of reads: blesbok, water buffalo, and white-tailed deer. All such dubious cases were based on matches either to the D-loop region of the control region, or to a conserved section of the 16S rDNA gene. All questionable reads resulted in perfect matches to other mammal species, including several ungulates known to occur in, and in the vicinity of, Mbuluzi (blue wildebeest, waterbuck, common duiker and domestic goat), when they were individually used in BLAST searches against GenBank. Whilst the control region as a whole is variable, its central conserved domain is one of the most conserved regions of the mitochondrial genome (Brown et al 1986), explaining the observed matches to multiple species of ungulates across the ~150 bp long matching reads. Therefore, for practical implementation of this method, we recommend disregarding any inference from matches to highly conserved mitochondrial sites.

The dung beetle gut approach appears to be at least as effective as using other invertebrates as a mammalian DNA source. Calvignac-Spencer et al. (2013b) extracted DNA from 201 flesh-feeding carrion flies in Côte d’Ivoire and Madagascar, and were able to detect a total of 26 mammal species, whereas Schnell et al. (2012) detected six mammal species from 25 leeches. It is, however, important to point out that the above two studies incorporated PCR-based amplification of targeted genes, whereas our approach circumvents the often laborious, costly, and time-intensive process of PCR optimisation and sequencing. Even more importantly, it removes the inherent bias-prone nature of differential primer-binding success of PCR, which is of particular concern when attempting to amplify sequences from multiple species of an undetermined, and genetically diverse fauna.

An obvious limitation of virtually all metagenomics studies is their reliance upon reliably identified, annotated, and curated reference sequences on publicly accessible databases for sequence assignment/identification. In the present study, this is unlikely to have been a major hindrance, because mitochondrial genome sequences were available for most of the ungulates likely to have been sources of dung from the sampling site. However, it will undoubtedly be a limitation if this technique is employed in areas where the bulk of the mammal fauna has not yet been thoroughly barcoded or sequenced. Because of this, and because of the present and projected rise in the number of metagenomics studies and biodiversity assessments, we support increasing the number of reliably identified reference sequences in public databases, including not only barcodes but also complete mitogenomes, which can now be reliably and economically obtained in bulk (Gillett et al. 2014; Crampton-Platt et al. 2015).

Whilst only a very small fraction of the total sequence reads generated matched mammalian sequences (2.0 × 10^−6^% matching mitogenomes, and 3.3 × 10^−6^% matching BOLD sequences), we were nevertheless able to assemble the near-complete mitogenome of the blue wildebeest from four of the samples, and this despite an almost undetectable initial DNA extraction concentration in one sample (DB009) prior to sequencing. That such long sequences can be successfully assembled increases the confidence of subsequent identifications, and further justifies the suggestion that impetus should be placed in increasing representation of mitogenomes in public databases.

One unexplored avenue, beyond the scope of this article, is the use of the nuclear DNA reads, perhaps through low coverage genome skimming (Dodsworth 2015) to further identify additional mammalian sequences. Whilst we believe that this is important, representation of nuclear sequences with species-level identifications on public databases is less comprehensive than that for mitochondrial sequences, which have traditionally been used for species-level assignment, especially through the DNA “barcode” (Hebert et al. 2002).

Although dung beetles have been put forward as a suitable proxy for mammal diversity, a key outstanding question is precisely how closely their diversities and abundances are correlated. Recent advances have enabled 100-fold enrichment of mtDNA prior to sequencing through use of a gene capture chip, with little detectable bias (Liu et al. 2016). Such technology could allow a similar eDNA metagenomics methodology as described here to be employed on a much larger scale, and with greater sequencing depth, allowing the sequencing of DNA pools from hundreds or thousands of beetle gut extractions sampled from different areas of known mammalian fauna.

In conclusion, we have demonstrated that mammalian DNA can be successfully sequenced and identified from direct sequencing of dung beetle intestinal content DNA extractions, circumventing the need for both direct observation of mammals, and laborious and bias-prone PCR reactions.

This methodology has the potential to be useful wherever rapid terrestrial mammalian biodiversity measurement might be necessary and dung beetles occur, and offers a novel approach to potentially detect and identify or monitor very rare and enigmatic mammals, or even those presumed to be locally or globally extinct. Examples of mammal populations which have proven to be exceptionally difficult to detect or monitor through traditional means, and which might potentially be ‘re-found’ using this methodology include those of Arabian Tahr (*Hemitragus jayakari*) and Arabian leopard (*Panthera pardus nimr*) supposedly still present in the mountains of the United Arab Emirates and Oman (Cunningham 2001; Edmonds 2006).

## Acknowledgments

We are grateful for the support of the British Ecological Society for awarding CG a small research grant (5524 / 6568) to undertake this study. We also acknowledge the assistance of the staff of the Mbuluzi Game Reserve and All Out Africa, for permission to sample at the site, and especially Morgan Vance and Kim Roques, for their logistical support of our fieldwork. We appreciated the help of many undergraduate students from the University of East Anglia field-course to Swaziland, who assisted CG in the field, especially Rebecca Easter, Elliot Reynolds, and Samantha Evans. We thank David Moraga of the University of Florida Interdisciplinary Center for Biotechnology Research for his assistance in preparing the samples for sequencing. Finally, we are indebted to Lewis Spurgin and Douglas Yu for their constructive criticism of the original grant application and an early draft of this article, and to Matthew Gage (all University of East Anglia) for his support of the project during its inception. None of the authors has a conflict of interest to declare.

## Data accessibility

Data resulting from this study, and examples of computer scripts used in its analysis, will be deposited in Dryad, once the manuscript is peer-reviewed and published.

## Figure and table legends

Table 1 - Summary of the number of matches to GenBank mammal mitogenome sequences from each of the ten dung beetle gut extractions, and the pooled sample.

Table 2 - Summary of the longest *de novo* mitogenome assembly contigs obtained for each of the dung beetle gut extractions and their taxonomic assignments using BLAST.

Tabel 3 - The number of R1 or R2 reads, for each dung beetle gut extraction sample, matching to a mammalian sequence in the BOLD database with 98% or greater identity over at least 100 bp.

Figure 1 - Schematic workflow of steps undertaken in the dung beetle gut DNA extraction metagenomics analysis.

## Supporting information

Appendix S1–Checklist of wild mammals recorded from Mbuluzi Game Reserve, Swaziland (PDF).

Appendix S2–The 25 mammal mitogenomes retrieved from GenBank, and used in sequencing read identification (PDF).

Appendix S3–The 25 mammal mitogenome sequences retrieved from GenBank, and used in the mitogenomic search (FASTA).

Appendix S4–The mammal barcode sequences retrieved from BOLD, and used in the barcode search (FASTA).

Appendix S5–Dung beetle sample identifications, and corresponding DNA extraction concentrations and number of sequence reads generated for each of the 10 species and samples selected for intestinal content DNA extraction, and the pooled dung sample (PDF).

Appendix S6–Results of the raw read quality control undertaken with Trimmomatic (PDF).

## Author contributions

CG and IB conceived the ideas and designed methodology; CG collected the data; CG and AJ analysed the data; CG and JH led the writing of the manuscript. All authors contributed critically to the drafts and gave final approval for publication.

## Appendix S1

Wild mammals recorded from the Mbuluzi Game Reserve, Lubombo, Swaziland.

**Macroscelididae (Elephant shrews)**

***Elephantulus brachyrhynchus***

(Short-snouted elephant-shrew)

**Soricidae (Shrews)**

***Crocidura gracilipes***

(Peters' musk shrew)

***Crocidura bicolor***

(Tiny musk shrew)

***Suncus lixus***

(Greater dwarf shrew)

***Crocidura flavescens***

(Greater musk shrew)

Swaziland Red Data Status: Least Concern

Southern African Red Data Status: Not listed

International Red Data Status: Vulnerable

***Crocidura cyanea***

(Reddish grey musk shrew)

***Crocidura hirta***

(Lesser red musk shrew)

**Chrysochloridae (Golden moles)**

***Amblysomus hottentotus***

(Hottentot's golden mole)

Southern African Red Data Status: Southern African endemic

**Pteropodidae (Fruit bats)**

***Rousettus aegyptiacus***

(Egyptian fruit bat)

***Epomophones wahlbergi***

(Wahlberg's epauletted fruit bat)

***Epomophorus crypturus***

(Peters' epauletted fruit bat)

**Emballonuridae (Sheath-tailed bats)**

***Taphozous mauritianus***

(Tomb bat)

**Molossidae (Free tailed bats)**

***Tadarida condylura***

(Angola free-tailed bat)

***Tadarida pumila***

(Little free-tailed bat)

***Tadarida aegyptiaca***

(Egyptian free tailed bat)

**Vespertilionidae (Vesper bats) Miniopterinae**

***Scotophilus viridis***

(Lesser yellow house bat)

***Eptesicus zuluensis***

(Aloe serotine bat)

***Chalinolobis variegatus***

(Butterfly bat)

***Miniopterus schreibersii***

(Schreibers' long fingered bat)

Swaziland Red Data Status: Near Threatened Southern

African Red Data Status: Not listed

International Red Data Status: Near Threatened

***Nycticeius schlieffenii***

(Schlieffen's bat)

**Vespertilionidae (Vesper bats) Vespertilioninae**

***Myotis tricolor***

(Temminck's hairy bat)

***Pipistrellus nanus***

(Banana bat)

***Pipistrellus kuhlii***

(Kuhl's bat)

***Eptesicus capensis***

(Cape serotine bat)

**Nycteridae (Slit faced bats)**

***Nycteris thebaica***

(Common slit faced bat)

**Rhinolophidae (Horseshoe bats)**

***Rhinolophus clivorus***

(Geoffroy's horseshoe bat)

***Rhinolophus darlingi***

(Darling's horseshoe bat)

***Rhinolophus simulator***

(Bushveld horseshoe bat)

**Hipposideridae (Trident and leaf-nosed bats)**

***Hipposideros caffer***

(Sundevall's leaf-nosed bat)

**Cercopithecidae (Monkeys and baboons)**

***Cercopithecus mitis***

(Samango monkey)

Swaziland Red Data Status: Endangered

Southern African Red Data Status: Rare

International Red Data Status: Not listed

***Papio ursinus***

(Chacma baboon)

(SiSwati: Imfene)

***Cercopithecus aethiops pygerythrus***

(Vervet monkey)

(SiSwati: Ingobiyane)

**Lorisidae (Bushbabies) Galaginae**

***Galago senegalensis***

(Lesser bushbaby)

***Galago crassicaudatus***

(Thick tailed bushbaby)

**Manidae (Pangolin)**

***Manis temminckii***

(Pangolin)

Swaziland Red Data Status: Endangered

Southern African Red Data Status: Vulnerable

International Red Data Status: Near Threatened

**Leporidae (Hares and rabbits)**

***Pronolagus crassicaudatus***

(Natal red rock rabbit)

***Lepus saxatilis***

(Scrub hare)

(SiSwati: Logwaja)

**Hystricidae (Porcupines)**

***Hystrix africaeaustralis***

(Porcupine)

**Gliridae (Dormice)**

***Graphiurus murinus***

(Woodland dormouse)

**Thryonomyidae (Canerats)**

***Thryonomys swinderianus***

(Greater canerat)

**Cricetidae and Muridae (Rats and mice) Otomyinae**

***Otomys angoniensis***

(Angoni vlei rat)

**Cricetidae and Muridae (Rats and mice) Murinae**

***Thamnomys dolichurus***

(Woodland mouse)

***Thamnomys cometes***

(Mozambique woodland mouse)

***Aethomys namaquensis***

(Namaqua rock mouse)

***Aethomys chrysophilus***

(Red veld rat)

***Dasymys incomtus***

(Water rat)

Swaziland Red Data Status: Vulnerable

Southern African Red Data Status: Indeterminate

International Red Data Status: Data Deficient

***Lemniscomys rosalia***

(Single striped mouse)

***Mus minutoides***

(Pygmy mouse)

***Mastomys natalensis***

(Natal multimammate mouse)

***Rattus rattus***

(House rat)

**Exotic**

***Thallomys paedulcus***

(Tree mouse)

**Cricetidae and Muridae (Rats and mice) Gerbillinae**

***Tatera leucogaster***

(Bushveld gerbil)

**Cricetidae and Muridae (Rats and mice) Cricetinae**

***Saccostomus campestris***

(Pouched mouse)

**Cricetidae and Muridae (Rats and mice) Dendromurinae**

***Dendromus melanotis***

(Grey climbing mouse)

***Dendromus mystacalis***

(Chestnut climbing mouse)

***Steatomys pratensis***

(Fat mouse)

**Hyaenidae (Aardwolf and hyaenas)**

***Proteles cristatus***

(Aardwolf)

(SiSwati: Singce)

Swaziland Red Data Status: Near Threatened

Southern African Red Data Status: Not listed

International Red Data Status: Not listed

***Crocuta crocuta***

(Spotted hyaena)

Swaziland Red Data Status: Vulnerable

Southern African Red Data Status: Not listed

International Red Data Status: Conservation-dependent

**Felidae (Cats)**

***Acinonyx jubatus***

(Cheetah)

Swaziland Red Data Status: Regionally Extinct

Southern African Red Data Status: Out of Danger

International Red Data Status: Vulnerable

***Panthera pardus***

(Leopard)

(SiSwati: Ingwe)

Swaziland Red Data Status: Near Threatened

Southern African Red Data Status: Rare

International Red Data Status: Not listed

***Felis serval***

(Serval)

(SiSwati: Indloti)

Swaziland Red Data Status: Near Threatened

Southern African Red Data Status: Rare

International Red Data Status: Not listed

***Felis lybica***

(African wild cat)

Swaziland Red Data Status: Data Deficient

Southern African Red Data Status: Vulnerable

International Red Data Status: Not listed

**Canidae (Foxes, wild dog and jackals)**

***Lycaon pictus***

(African wild dog)

Swaziland Red Data Status: Regionally Extinct

Southern African Red Data Status: Endangered

International Red Data Status: Endangered

***Canis adustus***

(Side-striped jackal)

***Canis mesomelas***

(Blackbacked jackal)

(SiSwati: Jakalazi)

**Mustelidae (Otters, polecats, weasels, honey badger)**

***Mellivora capensis***

(Honey badger)

Swaziland Red Data Status: Vulnerable

Southern African Red Data Status: Vulnerable

International Red Data Status: Not listed

***Ictonyx striatus***

(Striped polecat)

***Poecilogale albinucha***

(Striped weasel)

Swaziland Red Data Status: Near Threatened

Southern African Red Data Status: Rare

International Red Data Status: Not listed

**Viverridae (Mongooses, civets, genets and suricate)**

***Civettictis civetta***

(African civet)

Swaziland Red Data Status: Near Threatened

Southern African Red Data Status: Rare

International Red Data Status: Not listed

***Genetta tigrina***

(Large spotted genet)

***Herpestes ichneumon***

(Large gray mongoose)

***Galerella sanguinea***

(Slender mongoose)

***Rhynchogale melleri***

(Meller's mongoose)

Swaziland Red Data Status: Data Deficient

Southern African Red Data Status: Vulnerable

International Red Data Status: Not listed

***Ichneumia albicauda***

(White tailed mongoose)

***Atilax paludinosus***

(Water mongoose)

***Mungos mungo***

(Banded mongoose)

***Helogale parvula***

(Dwarf mongoose)

**Orycteropodidae (Antbear)**

***Orycteropus afer***

(Antbear)

**Equidae (Zebras)**

***Equus burchelli antiquorus***

(Burchells zebra)

(SiSwati: Lidvuba)

Swaziland Red Data Status: Least Concern

Southern African Red Data Status: Not listed

International Red Data Status: Not listed

**Suidae (Pigs)**

***Potamochoerus porcus***

(Bush pig)

(SiSwati: Ingulube)

***Phacochoerus aethiopicus***

(Warthog)

(SiSwati: Budzayikatane)

Swaziland Red Data Status: Least Concern

Southern African Red Data Status: Not listed

International Red Data Status: Not listed

**Bovidae (Buffalo, wildebeest and buck) Alcelaphinae (Wildebeest, hartebeest and blesbok)**

***Connochaetes taurinus***

(Blue wildebeest)

(SiSwati: Ingongoni)

Swaziland Red Data Status: Least Concern

Southern African Red Data Status: Not listed

International Red Data Status: Conservation-dependent

**Bovidae (Buffalo, wildebeest and buck) Cephalophinae (Duikers)**

***Cephalophus natalensis***

(Red duiker)

(SiSwati: Umsumphe)

Swaziland Red Data Status: Near Threatened

Southern African Red Data Status: Rare

International Red Data Status: Conservation-dependent

***Sylvicapra grimmia***

(Grey duiker)

(SiSwati: Impunzi)

**Bovidae (Buffalo, wildebeest and buck) Antilopinae (Klipspringer, dik dik, oribi, grysbok)**

***Oreotragus oreotragus***

(Klipspringer)

(SiSwati: Logoga)

Swaziland Red Data Status: Near Threatened

Southern African Red Data Status: Not listed

International Red Data Status: Conservation-dependent

***Ourebia ourebi***

(Oribi)

(SiSwati: Liwula)

Swaziland Red Data Status: Vulnerable

Southern African Red Data Status: Vulnerable

International Red Data Status: Conservation-dependent

***Raphicerus campestris***

(Steenbok)

***Raphicerus sharpei***

(Sharpe's grysbok)

**Bovidae (Buffalo, wildebeest and buck)**

**Aepycerotinae (Impala)**

***Aepyceros melampus***

(Impala)

(SiSwati: Imphala)

Swaziland Red Data Status: Least Concern

Southern African Red Data Status: Not listed

International Red Data Status: Conservation-dependent

**Bovidae (Buffalo, wildebeest and buck) Bovinae (Buffalo, kudu, bushbuck)**

***Tragelaphus angasi***

(Nyala)

Swaziland Red Data Status: Least Concern

Southern African Red Data Status: Not listed

International Red Data Status: Conservation-dependent

***Tragelaphus scriptus***

(Bushbuck)

Swaziland Red Data Status: Least Concern

Southern African Red Data Status: Not listed

International Red Data Status: Not listed

***Tragelaphus strepsiceros***

(Greater kudu)

Swaziland Red Data Status: Least Concern

Southern African Red Data Status: Not listed

International Red Data Status: Conservation-dependent

**Bovidae (Buffalo, wildebeest and buck) Reduncinae (Reedbuck, waterbuck, lechwe)**

***Redunca arundinum***

(Common reedbuck)

Swaziland Red Data Status: Near Threatened

Southern African Red Data Status: Not listed

International Red Data Status: Conservation-dependent

***Redunca fulvorufula***

(Mountain reedbuck)

(SiSwati: Lincala)

Swaziland Red Data Status: Near Threatened

Southern African Red Data Status: Not listed

International Red Data Status: Conservation-dependent

***Kobus ellipsiprymnus***

(Common waterbuck)

Swaziland Red Data Status: Near Threatened

Southern African Red Data Status: Not listed

International Red Data Status: Conservation-dependent

## Appendix S2

The 25 mammal mitogenomes retrieved from GenBank used in sequencing read identification.

| Common name | Scientific name | GenBank accession code |
| --- | --- | --- |
| Vervet monkey* | *Chlorocebus pygerythrus* | EF597500 |
| Domestic cattle | *Bos taurus* | EU177832 |
| Blesbok | *Damaliscus pygargus* | FJ207530 |
| Domestic goat | *Capra hircus* | GU295658 |
| Impala* | *Aepyceros melampus* | JN632592 |
| Blue wildebeest* | *Connochaetes taurinus* | JN632627 |
| Waterbuck* | *Kobus ellipsiprymnus* | JN632651 |
| Nyala* | *Tragelaphus angasii* | JN632702 |
| Greater Kudu* | *Tragelaphus strepsiceros* | JN632708 |
| Cape baboon* | *Papio ursinus* | JX946204 |
| Weasel | *Mustela nivalis* | KT693382 |
| Hippopotamus | *Hippopotamus amphibius* | NC_000889 |
| African bush elephant | *Loxodonta africana* | NC_000934 |
| Plains Zebra* | *Equus quagga* | JX312733 |
| Domestic horse | *Equus caballus* | NC_001640 |
| Domestic cat | *Felis catus* | NC_001700 |
| Common wallaroo | *Macropus robustus* | NC_001794 |
| House mouse | *Mus musculus* | NC_005089 |
| Common warthog* | *Phacochoerus africanus* | NC_008830 |
| Giraffe* | *Giraffa camelopardalis* | NC_012100 |
| Human | *Homo sapiens* | NC_012920 |
| African palm civet | *Nandinia binotata* | NC_024567 |
| Greater mouse-eared bat | *Myotis myotis* | NC_029346 |
| Domestic dog | *Canis familiaris* | U96639 |
| Northern white rhinoceros | *Ceratotherium simum* | Y07726 |

\*Wild mammals occurring in Mbuluzi Game Reserve, Swaziland.

## Appendix S5

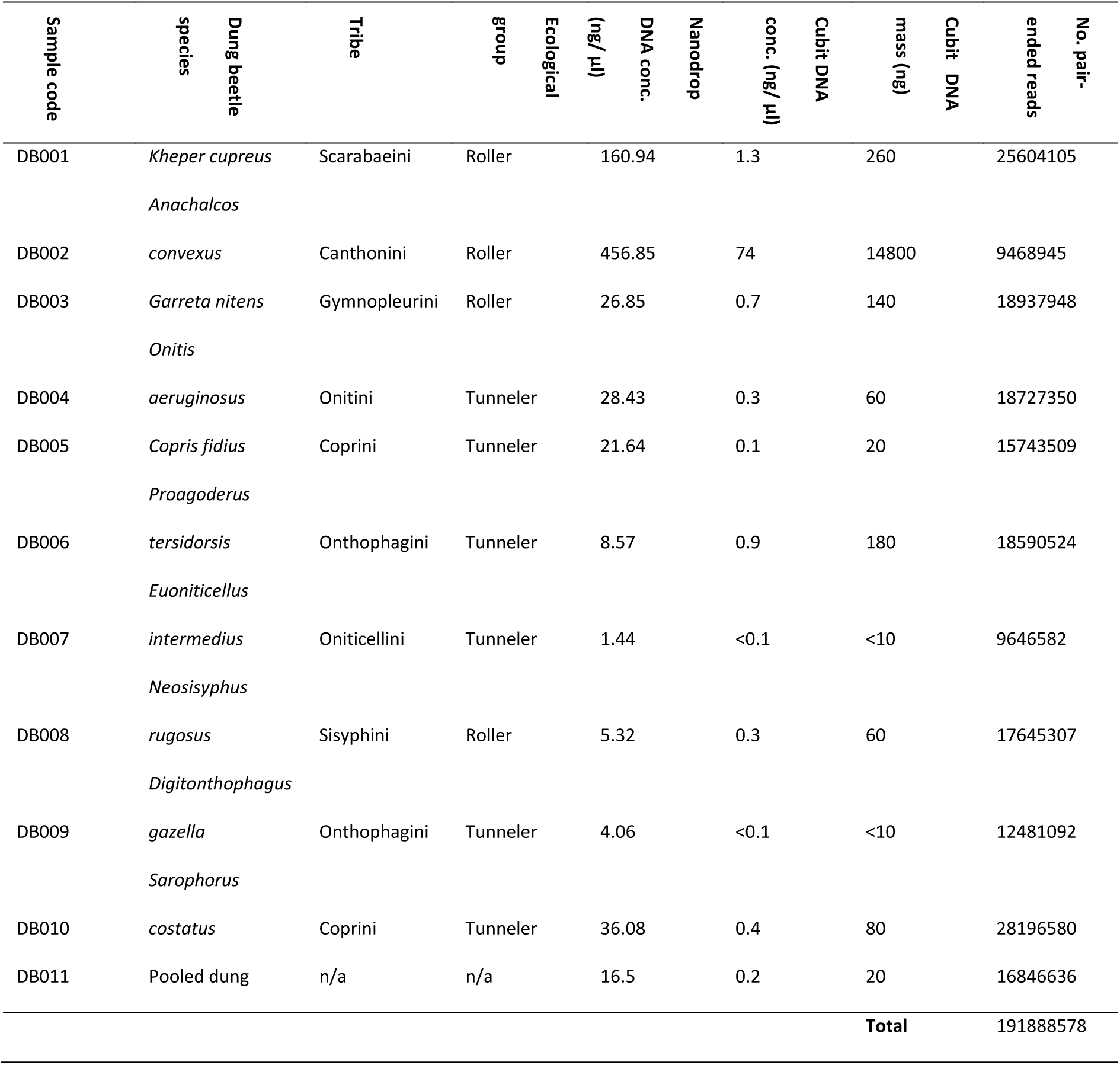
Dung beetle sample identifications, and corresponding DNA extraction concentrations and number of sequence reads generated for each of the 10 species selected for intestinal content DNA extraction, and the pooled dung sample.

## Appendix S6

Results of the raw read quality control undertaken with Trimmomatic (PDF).

Appendix S6 Results of the raw read quality control undertaken with Trimmomatic (PDF).
| Sample | Input<br>read pairs | Both<br>surviving | Forward<br>only<br>surviving | Reverse<br>only<br>surviving | Dropped | Both<br>surviving<br>% | Forward<br>only<br>surviving | Reverse<br>only<br>surviving | Dropped<br>% |
| --- | --- | --- | --- | --- | --- | --- | --- | --- | --- |
| DB001 | 25604105 | 25459419 | 28547 | 76697 | 39442 | 99.43 | 0.11 | 0.3 | 0.15 |
| DB002 | 9468945 | 9369837 | 9255 | 11743 | 78110 | 98.95 | 0.1 | 0.12 | 0.82 |
| DB003 | 18937948 | 18914107 | 16868 | 3733 | 3240 | 99.87 | 0.09 | 0.02 | 0.02 |
| DB004 | 18727350 | 17221800 | 15885 | 1477255 | 12410 | 91.96 | 0.08 | 7.89 | 0.07 |
| DB005 | 15743509 | 13640961 | 11981 | 2081896 | 8671 | 86.64 | 0.08 | 13.22 | 0.06 |
| DB006 | 18590524 | 18567521 | 17273 | 3258 | 2472 | 99.88 | 0.09 | 0.02 | 0.01 |
| DB007 | 9646582 | 8450155 | 7803 | 1178249 | 10375 | 87.6 | 0.08 | 12.21 | 0.11 |
| DB008 | 17645307 | 14637551 | 13733 | 2964988 | 29035 | 82.95 | 0.08 | 16.8 | 0.16 |
| DB009 | 12481092 | 10493771 | 10264 | 1961741 | 15316 | 84.08 | 0.08 | 15.72 | 0.12 |
| DB010 | 28196580 | 22303763 | 21633 | 5766016 | 105168 | 79.1 | 0.08 | 20.45 | 0.37 |
| DB011 | 16846636 | 14479625 | 13565 | 2328192 | 25254 | 85.95 | 0.08 | 13.82 | 0.15 |
| Total paired reads | 191888578 | 173538510 | 166807 | 17853768 | 329493 |  |  |  |  |
| Total reads | 383777156 | 347077020 |  |  |  |  |  |  |  |

## References

Altschul, S.F., Gish, W., Miller, W., Myers, E.W. & Lipman, D.J. (1990) Basic local alignment search tool. Journal of Molecular Biology, 215, 403–410.

Barnett, A. (1995) Expedition field techniques: Primates. Royal Geographical Society, London.

Barnett, A. & Dutton, J. (1995) Expedition field techniques: Small mammals. Royal Geographical Society, London.

Beng, K.C., Tomlinson, K.W., Shen, X.H., Surget-Groba, Y., Hughes, A.C., Corlett, R.T., & Slik, J.W.F. (2016) The utility of DNA metabarcoding for studying the response of arthropod diversity and composition to land-use change in the tropics. Scientific Reports, 6, 1–13.

Benson, D.A., Cavanaugh, M., Clark, K., Karsch-Mizrachi, I., Lipman, D.J., Ostell, J. & Sayers, E.W. (2013) GenBank. Nucleic Acids Research 41: D36–42.

Bohmann, K., Evans, A., Gilbert, M.T.P., Carvalho, G.R., Creer, S., Knapp, M., Yu, D.W. & de Bruyn, M. (2014) Environmental DNA for wildlife biology and biodiversity monitoring. Trends in Ecology and Evolution, 29, 358–367.

Bolger, A.M., Lohse, M. & Usadel, B. (2014) Trimmomatic: a flexible trimmer for Illumina sequence data. Bioinformatics, 30, 2114–2120.

Brown, G. (1986) Structural conservation and variation in the D-loop-containing region of vertebrate mitochondrial DNA. Journal of Molecular Biology, 192, 503–511.

Calvignac-Spencer, S., Leedndertz, F.H., Gilbert, M.T.P. & Schubert, G. (2013a) An invertebrate stomach’s view on vertebrate ecology. Bioessays, 35, 1004–1013.

Calvignac-Spencer, S., Merkel, K., Kutzner, N., Kühl, H., Boesch, C., Kappeler P.M., Metzger, S., Schubert, G. & Leendertz, F.H. (2013b) Carrion fly-derived DNA as a tool for comprehensive and cost-effective assessment of mammalian biodiversity. Molecular Ecology, 22, 915–924.

Cambefort, Y. (1991) Biogeography and evolution. Dung Beetle Ecology (eds I. Hanski & Y. Cambefort), pp. 51–67. Princeton University Press, Princeton.

Cambefort, Y. & Hanski, I. (1991) Dung beetle population biology. Dung Beetle Ecology (eds I. Hanski & Y. Cambefort), pp. 36–50. Princeton University Press, Princeton.

Corlett, R.T. (2016) A Bigger Toolbox: Biotechnology in Biodiversity Conservation. Trends in Biotechnology.

Crampton-Platt, A., Yu, D.W., Zhou, X. & Vogler, A.P. (2016) Mitochondrial metagenomics: letting the genes out of the bottle. GigaScience, 5, 1–11.

Cunningham, P.L. (2001) On the distribution and status of the Arabian Tahr, *Hemitragus jayakari*, in the United Arab Emirates and northern Oman. Zoology in the Middle East, 23, 13–16.

Creer, S., Deiner, K., Frey, S., Porazinska, D., Taberlet, P., Thomas, W.K., Potter, C. & Bik, H.M. (2016) The ecologist’s field guide to sequence-based identification of biodiversity. Methods in Ecology and Evolution, doi: 10.1111/2041-210X.12574.

Davis, A. L. V., Frolov, A. V. & Scholtz., C. H. (2008) The African dung beetle genera. Protea Book House, Pretoria.

Deschodt, C.M., Davis, L.V. & Scholtz, C.H. (2015) A new synonymy in the *fidius* group of *Copris* Müller 1764 (Coleoptera: Scarabaeidae: Scarabaeinae) and a new species from the highland grasslands of South Africa. Zootaxa, 3949, 431–438.

Dodsworth, S. (2015) Genome skimming for next-generation biodiversity analysis. Trends in Plant Science, 20, 525–527.

Drummond, A.J., Newcomb, R.D., Buckley, T.R., Xie, D. & Dopheide, A. Evaluating a multigene environmental DNA approach for biodiversity assessment. GigaScience, 4, 1–20.

Edgar, R.C. (2010) Search and clustering orders of magnitude faster than BLAST. Bioinformatics, 26, 2460–2461.

Edmonds, J-A, Budd, K.J., al Midfa, A. & Gross, C. (2006) Status of the Arabian leopard in United Arab Emirates. Cat News, 1, 33–39.

Ferreira, M.C. (1972) Os escarabideos de Africa (sul do Saara) I. Revista de Entomologia de Moçambique, 11, 1–1088.

Gardner, T.A., Barlow, J., Araujo, I.S., Avila-Pires, T.C., Bonaldo, A.B., Costa, J.E., Esposito, M.C., Ferreira, L.V., Hawes, J., Hernandez, M.I.M., et al. (2008) The cost-effectiveness of biodiversity surveys in tropical forests. Ecology Letters, 11, 139–150.

Gillett, C.P.D.T., Crampton-Platt, A., Timmermans, M.J.T.N., Jordal, B.J., Emerson, B.C. & Vogler, A.P. (2014) Bulk de novo mitogenome assembly from pooled total DNA elucidates the phylogeny of weevils (Coleoptera: Curculionoidea). Molecular Biology and Evolution, 31, 2223–2237.

Halffter, G. & Favila, M.E. (1993) The Scarabaeinae (Insecta: Coleoptera) an animal group for analysing, inventorying and monitoring biodiversity in tropical rainforest and modified landscapes. Biology International, 27, 15–21.

Halffter., G. & Matthews, E.G. (1966) The natural history of dung beetles of the subfamily Scarabaeinae (Coleoptera: Scarabaeidae). Folia Entomologica Mexicana, 12, 1–312.

Hanski, I. & Cambefort, Y. (Eds.) (1991) Dung Beetle Ecology. Princeton University Press, Princeton.

Hebert, P.D.N., Cywinska A., Ball, S.L. & DeWaard, J.R. (2003) Biological identifications through DNA barcodes. Proceedings of the Royal Society B-Biological Sciences, 270, 313–321.

Humphries, C.J., Williams, P.H. & Vane-Wright, R.I. (1995) Measuring biodiversity value for conservation. Annual Review of Ecology, Evolution, and Systematics, 26, 93–111.

Kearse, M., Moir, R., Wilson, A., Stones-Havas, S., Cheung, M., Sturrock, S., Buxton, S., Cooper, A., Markowitz, S., Duran, C., et al. (2012) Geneious Basic: An integrated and extendable desktop software platform for the organization and analysis of sequence data. Bioinformatics, 28, 1647–1649.

Krebs, C.J. (2006) Mammals. Ecological Census Techniques, 2^nd^ edn. (ed W.J. Sutherland), 351–369. Cambridge University Press, Cambridge.

Larsen, T.H., Lopera, A. & Forsyth, A. (2006) Extreme trophic and habitat specialization by Peruvian dung beetles (Coleoptera: Scarabaeidae: Scarabaeinae). Coleopterists Bulletin, 60, 315–324.

Larsen, T.H. (2011) Dung beetles of the Kwamalasamutu region, Suriname (Coleoptera: Scarabaeidae: Scarabaeinae). RAP Bulletin of Biological Assessment, 63, 91–103.

Liu, S., Wang, X., Xie, L., Tan, M., Li, Z., Su, X., Zhang, H., Misof, B., Kjer, K.M., Tang, M. & Niehuis, O., (2016) Mitochondrial capture enriches mito-DNA 100 fold, enabling PCR-free mitogenomics biodiversity analysis. Molecular Ecology Resources, 16, 470–479.

Mehus, J.O. & Vaughan, J. (2013) Molecular identification of vertebrate DNA and hemoparasite DNA within mosquito blood meals from eastern North Dakota. Vector-borne and Zoonotic Diseases, 13, 818–824.

Palestrini, C. (1992) Sistematica e zoogeografia del genere *Onthophagus* sottogenere *Proagoderus* Lansberge. Memorie della Società Entomologica Italiana, 124, 1–358.

Pokorny, S. & Zídek, J. (2015) Checklist of *Kheper* Janssens and description of a new species from northern Tanzania (Coleoptera: Scarabaeidae: Scarabaeinae). Insecta Mundi, 0429, 1–7.

Ratnasingham, S. & Hebert, P.D.N. (2007). BOLD: The Barcode of Life Data System (www.barcodinglife.org). Molecular Ecology Notes, 7, 355–364.

Reid, W.V., McNeely, J.A., Tunstal,l D.B., Bryant, D.A. & Winograd, M. (1993) Biodiversity Indicators for Policy-Makers. World Resources Institute, Washington, DC.

Royal Society (2003) Measuring biodiversity for conservation. Royal Society, London.

Schnell, I.B., Thomsen, P.F., Wilkinson, N., Rasmussen, M., Jensen, L.R.D., Willerslev, E., Bertelsen, M.F. & Gilbert, M.T.P. (2012) Screening mammal biodiversity using DNA from leeches. Current Biology, 22, 262–263.

Schipper, J., Chanson, J.S., Chiozza, F., Cox, N.A., Hoffmann, M., Katariya, V., Lamoreux, J., Rodrigues, A.S., Stuart, S.N., Temple, H.J. et al. 2008. The status of the world's land and marine mammals: diversity, threat, and knowledge. Science, 322, 225–230.

Spector, S. (2006) Scarabaeinae dung beetles (Coleoptera: Scarabaeidae: Scarabaeinae): an invertebrate focal taxon for biodiversity research and conservation. Coleopterists Bulletin, 60, 71–83.

Sutherland, W.J. (Ed.) (2006) Ecological Census Techniques, 2^nd^ edn. Cambridge University Press, Cambridge.

Townzen, J.S., Brower, A.V.Z. & Judd, D.D. (2008) Identification of mosquito bloodmeals using mitochondrial cytochrome oxidase subunit I and cytochrome b gene sequences. Medical and Vetenary Entomology, 22, 386–393.

Yang, C., Wang, X., Miller, J.A., de Blecourt, M., Ji, Y., Yang, C., Harrison, R.D., & Yu, D.W. (2014) Using metabarcoding to ask if easily collected soil and leaf-litter samples can be used as a general biodiversity indicator. Ecological Indicators 46, 379–389.

